# A systematic atlas of chaperome deregulation topologies across the human cancer landscape

**DOI:** 10.1101/122044

**Authors:** Ali Hadizadeh Esfahani, Angelina Sverchkova, Julio Saez-Rodriguez, Andreas A Schuppert, Marc Brehme

## Abstract

Proteome balance is safeguarded by the proteostasis network (PN), an intricately regulated network of conserved processes that evolved to maintain native function of the diverse ensemble of protein species, ensuring cellular and organismal health. Proteostasis imbalances and collapse are implicated in a spectrum of human diseases, from neurodegeneration to cancer. The characteristics of PN disease alterations however have not been assessed in a systematic way. Zooming in on the chaperome as a central PN component we turned to a curated functional ontology of the human chaperome that we connect in a high-confidence physical protein-protein interaction network. Challenged by the lack of a systems-level understanding of proteostasis alterations in the heterogeneous spectrum of human cancers, we assessed gene expression across more than 10,000 patient biopsies covering 22 solid cancers. We derived a novel customized Meta-PCA dimension reduction approach yielding M-scores as quantitative indicators of disease expression changes to condense the complexity of cancer transcriptomics datasets into quantitative functional network topographies. We confirm upregulation of the HSP90 family and also highlight HSP60s, Prefoldins, HSP100s, ER- and mitochondria-specific chaperones as pan-cancer enriched. Our analysis also reveals a surprisingly consistent strong downregulation of small heat shock proteins (sHSPs) and we stratify two cancer groups based on the preferential upregulation of ATP-dependent chaperones. Strikingly, our analysis highlight similarities between stem cell and cancer proteostasis, and diametrically opposed chaperome deregulation between cancers and neurodegenerative diseases. We developed a web-based Proteostasis Profiler tool (Pro^2^) enabling intuitive analysis and visual exploration of proteostasis disease alterations using gene expression data. Our study showcases a comprehensive profiling of chaperome shifts in human cancers and sets the stage for a systematic global analysis of PN alterations across the human diseasome towards novel hypotheses for therapeutic network re-adjustment in proteostasis disorders.

## Introduction

Eukaryotic proteomes comprise a complex repertoire of diverse protein species that are organized in a modular interactome network in order to execute native function in support of proteostasis and a healthy cellular phenotype. Proteome balance is safeguarded by the proteostasis network (PN), an intricately regulated network of conserved processes that have evolved to safeguard the healthy folded proteome (Balch et al. 2008). Cellular proteostasis capacity is limited within the constraints of each cell’s proteostasis boundary (Powers et al. 2009). Proteostasis imbalances, deficiency and functional collapse are implicated in a broad and increasing spectrum of protein conformational diseases with loss of native function or gain of toxic function, ranging from metabolic and neurodegenerative diseases to cancer (Hutt et al. 2009, Brehme et al. 2014). Increasing awareness of the fundamental role of the PN in cellular health, its relevance in diseases and potential as a therapeutic target of proteostasis regulator (PR) drugs call for a systematic and systems-level assessment of PN deregulation throughout the human diseasome, towards improved understanding of diseases of proteostasis deficiency and rationalized network-informed approaches to therapeutic proteostasis re-adjustment.

Important progress has been made in our understanding of proteostasis biology, building on fundamental insights on conserved proteostasis processes and their role in disease, such as chaperone-assisted protein folding and quality control (Hartl and Hayer-Hartl 2002, Young et al. 2004, Hartl et al. 2011, Kim et al. 2013), clearance through autophagy (Ohsumi 2001, Behrends et al. 2010, Janku et al. 2011, Rubinsztein et al. 2012, Türei et al. 2015) and the ubiquitin-proteasome system (UPS) (McNaught et al. 2001, Turnbull et al. 2007, Ciechanover 2015), followed by the appreciation of their concerted action within a conserved tightly regulated PN (Balch et al. 2008, Powers and Balch 2013). The identification, development and first clinical evaluations of small molecule PR drugs for therapeutic re-adjustment of proteostasis diseases such as cystic fibrosis represents a novel and powerful therapeutic paradigm (Balch et al. 2008, Powers et al. 2009, Bouchecareilh and Balch 2011, Calamini et al. 2011, Ramsey et al. 2011, Silva et al. 2011, Brandvold and Morimoto 2015, Das et al. 2015). First investigations have started to explore systems-level quantitative and functional approaches to assess the implications of PN functional arms such as the chaperome in human tissue aging and disease (Taipale et al. 2010, Brehme et al. 2014, Taipale et al. 2014, Rodina et al. 2016). A precise understanding of the molecular mechanisms by which PN alterations contribute to disease could open novel therapeutic intervention strategies in a wide spectrum of proteostasis-related diseases. Still, to date, there has been no systematic study addressing the characteristics and extent of PN alterations in human diseases at a systems-level.

The folding functional arm, the human chaperome, is highly conserved and of central importance in the PN, responsible for maintaining the native folded proteome. In cancers, mutations and genomic instability inevitably entail alterations of proteome composition and balance that are far less well explored than the consequences of nucleic acid sequence alterations. Post-translational alterations at the proteomic level are beyond the reach of DNA repair mechanisms and cancer cells are constantly challenged by the need to accommodate large amounts of proteotoxic stress in consequence of increased translational flux and proliferation as well as proteotoxic stressors. Proteome instability and pathological alterations in the abundance of key signalling or housekeeping molecules such as kinases, metabolic enzymes or molecular transporters have to be buffered by the PN to ensure cellular survival. The cancerous state poses characteristic requirements on the PN, such as high chaperone levels and elevated proteasome activity in order to ensure for sufficient correction or elimination of aberrant protein species in light of increased translational flux and metabolic stress (Whitesell and Lindquist 2005). This chronic challenge ultimately drives cancer cells into a dependency on quality control and stress response mechanisms, a phenomenon previously described as non-oncogene addiction (Dai et al. 2007, Solimini et al. 2007). Several individual chaperones and heat shock proteins such as HSP90 have consistently been found upregulated in cancers (Whitesell and Lindquist 2005). However, the profile and extent of chaperome differential expression has not been assessed systematically across the human cancer landscape.

Challenged by the genetic complexity and heterogeneity, collective prevalence and unmet medical need of the wide spectrum of human cancers as well as the lack of a systems-level understanding of proteostasis alterations during carcinogenic transformation, we developed a novel integrated analytical pipeline and software toolkit for the quantitative profiling of chaperome changes across the human cancer landscape (Fig 1). We utilized an expert-curated functional chaperome ontology comprising the ensemble of 332 human chaperone and co-chaperone genes (Brehme et al. 2014) (Fig 1A). In order to apply our analytical workflow on a recent and comprehensive cancer gene expression dataset with clinical relevance, we turned to The Cancer Genome Atlas (TCGA) compendium (Cancer Genome Atlas Research et al. 2013). We started with a customized genomic analysis pipeline in order to map chaperome functional family expression changes across TCGA solid cancers compared to matching normal tissue (Fig 1B). The resulting top-level view on cancer chaperome deregulation revealed a broad chaperome upregulation throughout the majority of cancers. This consistent and high overall chaperome upregulation prompted us to zoom in on functional sub-families. This analysis surfaced clusters of chaperome functional family up- and downregulation signatures that enabled further stratification of cancers. In summary, our analysis of the 10 major chaperome functional families reveals pronounced tissue differences of cancer chaperome deregulation. The preferential upregulation of ATP-dependent chaperone families such as HSP90s and HSP60s, while ATP-independent chaperones, co-chaperones, and small heat shock proteins (sHSPs) are consistently downregulated, is opposed to chaperome alteration patterns observed in brain tissues during aging and in neurodegenerative diseases (Brehme et al. 2014). These characteristic chaperome-wide differences further justify our approach and need for systematic maps of PN deregulation across the human diseasome.

**Fig 1.**
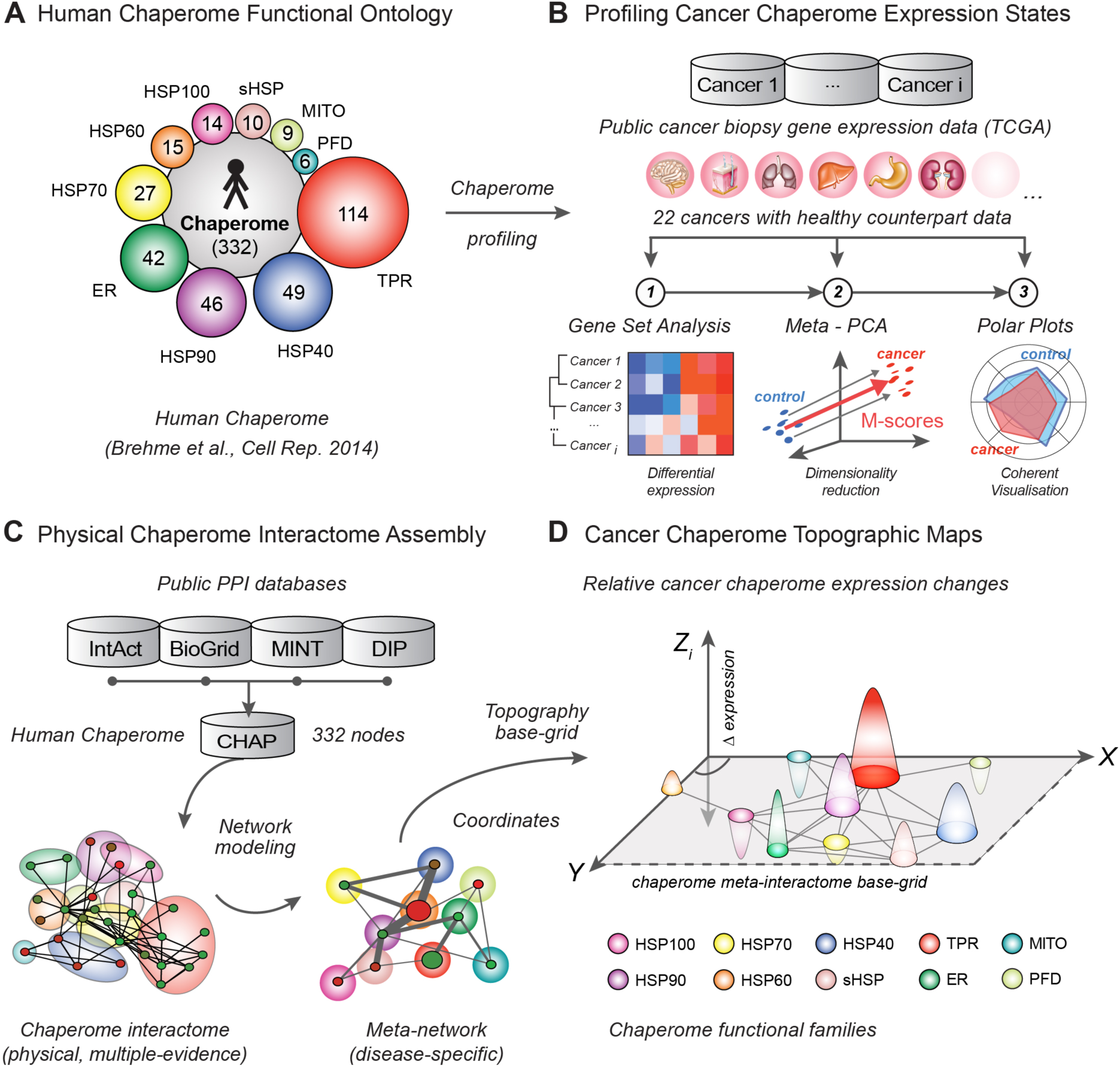
Chaperome Cancer Landscape Profiling. **A.** The human chaperome is a central PN functional arm in charge of maintaining the cellular folding environment. It comprises 332 chaperones and co-chaperones organized in 10 functional families (Brehme et al. 2014). **B.** Pipeline involving (1) Gene Set Analysis (GSA), (2) Meta-PCA, a novel two-step principal component analysis (PCA) - based dimension reduction approach yielding M-scores for quantitative analysis of chaperome changes across a compendium of TCGA solid cancer biopsy RNA-seq expression data, and (3) Polar Plots visualising contextual quantitative chaperome alterations. **C.** We connect 332 human chaperome genes (nodes) in a high-confidence literature-curated physical protein-protein interactome network (edges) and collapse nodes within functional families and edges between families into meta-nodes and meta-edges, respectively. The resulting optimized meta-networks serve as base-grid layout to enable interactome-guided chaperome landscape modeling. **D.** We use the high-confidence chaperome physical meta-interactome-guided base grid layout (X-Y dimensions) and Meta-PCA derived M-scores, indicating cancer expression change (Z dimension), to chart 3-dimensional quantitative topographic chaperome maps. Heatmaps, polar plots, meta-networks and 3D topographic map visualisations are accessible through the Proteostasis Profiler (Pro^2^) web-tool.

In order to enable comprehensive, contextual, and quantitative representations of the complexity of chaperome alterations across a large number of patient biopsy disease datasets, we developed a new custom data dimensionality reduction and visualisation approach. Combining Meta-PCA, a novel principal component analysis (PCA) based two-step dimension reduction algorithm and its resulting quantitative M-scores of chaperome functional family disease alteration of gene expression with contextual polar plot visualisations, we provide intuitive quantitative maps of cancer chaperome gene expression changes (Fig 1B).

The mechanistic understanding of genotype-phenotype relationships in complex genetically heterogeneous diseases such as cancers requires the consideration of the cellular interactome network (Sahni et al. 2013). To reduce complexity and to highlight contextual changes of chaperome functional families, we first generated a custom curated high-confidence physical protein-protein interaction (PPI) chaperome network. We then collapsed proteins (nodes) and physical PPIs (edges) into a meta-network, where meta-nodes represent respective functional family members and meta-edges bundle the interactions between families (Fig 1C). This curated high-quality chaperome meta-network base-grid enabled the contextual projection of cancer-specific chaperome functional family differential gene expression (M-scores) onto the underlying interactome. We integrated these dimensions into interactome-guided, three-dimensional topographic maps visualising chaperome functional family cancer differential gene expression changes in the context of interactome network proximity, intuitively providing quantitative views of cancer chaperome deregulation (Fig 1D).

To make these resources easily available to the community, we developed Proteostasis Profiler (Pro^2^), an integrated web-based suite of applications enabling intuitive quantitative analyses and comparative visualisation of differential expression of complex PN alterations across large disease dataset compendia such as the TCGA. Visualisation and analysis features include heat map clustering and polar plot display. Integrated meta-networks and interactome-guided 3D topographic maps ease comparative exploration of cancer chaperome deregulation in the context of interactome network wiring. Pro^2^ is designed to serve the scientific community as a user-friendly application for systems-level exploration of PN disease alterations, at reduced complexity.

Overall, this study represents a systematically derived systems-level atlas of chaperome deregulation maps in cancers and neurodegenerative diseases, with a detailed focus on chaperome functional family alterations. The integrated genomic analysis workflow, built into the Pro^2^ suite of visualisation tools, provides a resource and analytical platform for future characterisation and exploration of PN deregulation patterns across the human diseasome, and as a readout interface for network shifts induced by therapeutic regulation.

## Results

### Systematic Differential Gene Expression Profiling Highlights Functional Clusters of Chaperome Deregulation in Human Cancers

Homeostasis of the cellular proteome, or proteostasis, is fine-tuned by the proteostasis network (PN), an intricately regulated network of conserved processes that have evolved to safeguard the native functional proteome and cellular health. The human chaperome, an ensemble of 332 chaperones and co-chaperones, represents a central functional arm within the PN in charge of maintaining the cellular folding landscape (S1 Table A) (Brehme et al. 2014). Motivated by the genetic heterogeneity of cancers, their prevalence and associated medical need as well as the lack of a systems-level understanding of the role of proteostasis genomic alterations during carcinogenesis, we systematically assessed chaperome gene expression changes across the diverse spectrum of human cancers. We focused on an established resource of human cancer patient biopsy RNA-seq datasets provided through The Cancer Genome Atlas (TCGA) (Cancer Genome Atlas Research et al. 2013, Schubert et al. 2016). We considered 22 human solid cancers with available corresponding healthy counterpart tissue biopsy data. To obtain global views on chaperome commonalities or differences between cancers, we applied Gene Set Analysis (GSA) in order to quantify gene expression changes of the chaperome and its functional families. GSA is an advanced derivative of Gene Set Enrichment Analysis (GSEA) that methodologically differs primarily through its use of the maxmean statistic, the mean of the positive or negative gene scores in each gene set, whichever is larger in absolute value, that has proven superior to the modified Kolmogorov-Smirnov statistic used in GSEA (Efron and Tibshirani 2007). Secondly, GSA uses a different null distribution for false discovery rate (FDR) estimations, through a restandardiation of genes in addition to sample permutation in GSEA. This step is crucial, as it allows assessing statistical robustness of the expert-curated chaperome functional ontology gene family groups. We obtained the GSA derived probability (p values) for each functional gene group to be significantly up- or downregulated in cancer as ∆GSA values in the interval [-1, +1] according to ((1 - upregulation p value) - (1 - downregulation p value)).

Notably, the human chaperome is predominantly upregulated across the majority of TCGA solid cancers with a ∆GSA group mean change of +0,50 as compared to 100 random sets of non-chaperome genes (Fig 2A). This overall chaperome upregulation highlights cellular non-oncogene addiction to chaperone-assisted folding and protein quality control mechanisms in consequence of increased client load, further challenging cellular proteostasis and driving “proteostasis addiction” in cancers (Drummond and Wilke 2008). Despite the diverse established knowledge about the role of chaperone upregulation in cancer, the deregulation of the human chaperome has not been assessed at a systems-level throughout the human cancer landscape. To functionally resolve the general chaperome upregulation across cancers, we zoomed in on functional family gene expression alterations. GSA followed by Euclidean clustering of chaperome functional families revealed characteristic cancer differences. We found the key ATP-dependent HSP90 and HSP60 families, of which selected members have previously been shown to be upregulated in cancers, amongst the most highly upregulated functional families with ∆GSA group mean changes of +0.55 and +0.51, respectively, alongside ER-specific chaperone factors (+0,53), followed by Prefoldins (PFDs, +0,40), HSP100 AAA+ ATPases (+0,32), and mitochondria-specific chaperones (MITOs, +0,09) (Fig 2B). These six functional families of predominantly ATP-dependent chaperones represent an upregulation cluster with an overall group mean change of +0,40. Intriguingly, the HSP70-HSP40 system and the large family of TPR-domain containing co-chaperones are overall repressed, with less consistent and largely cancer-specific alterations. HSP40 co-chaperones (-0,09) cluster closest with HSP70s (-0,10), indicative of the functional relationship they engage in during the HSP70 chaperone cycle. While HSP40 co-chaperones are overall weakly downregulated (-0.09), also the second group of co-chaperones, the TPR-domain containing proteins, clustered with the HSP70-HSP40 system and were overall downregulated (-0,18). Strikingly, sHSPs (-0,74) were overall very consistently and strongly downregulated. Overall, sHSPs, TPRs, and the HSP70-HSP40 system clustered in a downregulation cluster with an overall group mean change of -0,28 across cancers.

**Fig 2.**
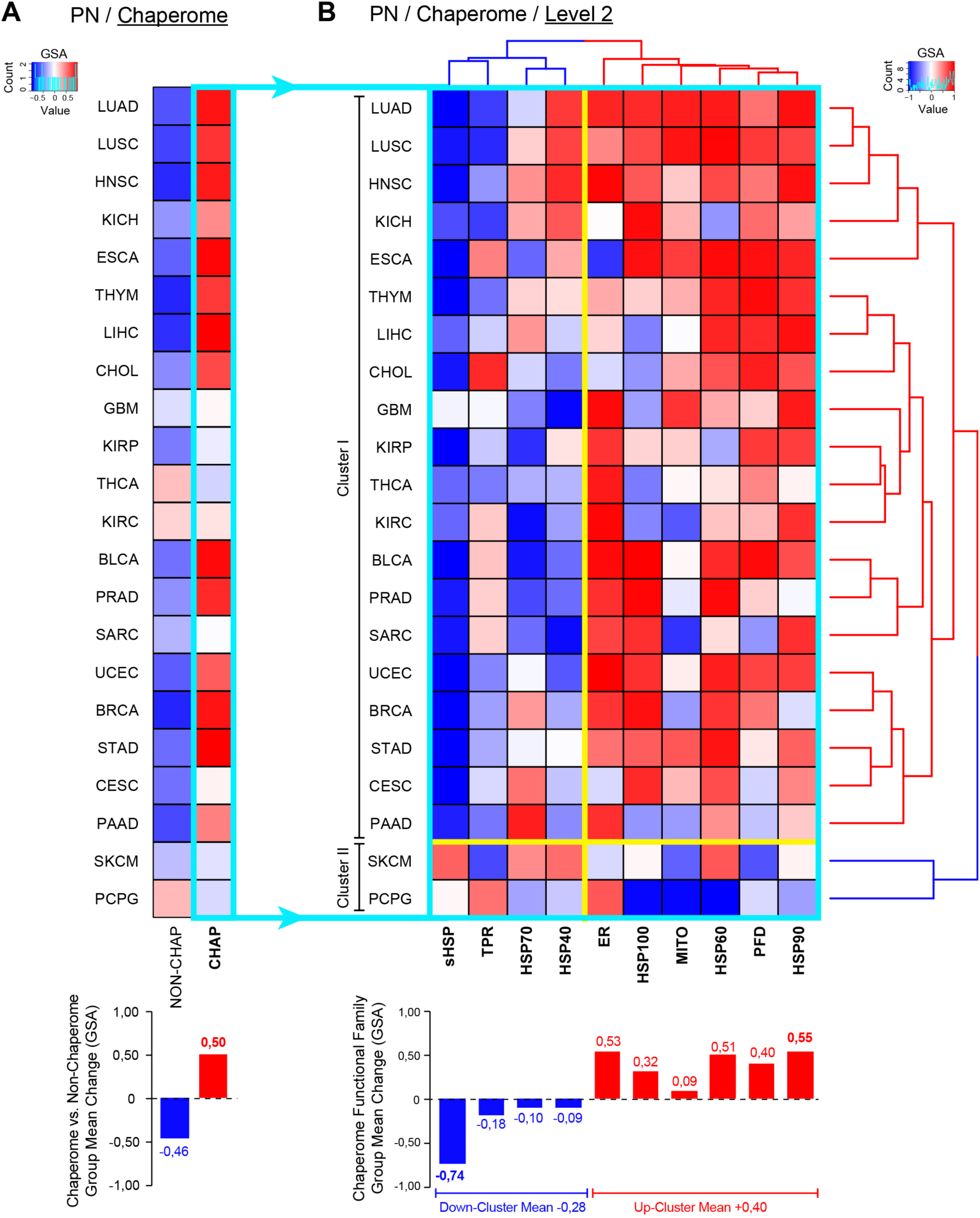
Chaperome Gene Expression Alterations in Human Cancers. Chaperome as compared to permutations of non-chaperome genes (**A**) and chaperome functional family (Level 2) (**B**) gene expression states in human cancer RNA-seq datasets from The Cancer Genome Atlas (TCGA) explored by Gene Set Analysis (GSA). Heatmaps indicate significance of up or down-regulation of cancer versus healthy gene expression as ∆GSA values in the interval [-1, +1], where ‘+1’ indicates significant upregulation (p value = 0), while ‘-1’ indicates significant downregulation (p value = 0). Chaperome functional families (B) are clustered by Euclidean distance (dendrograms). Bar graphs in A and B indicate functional family GSA group mean changes. Order of TCGA cancer groups (rows) in A is according to Euclidian distance of chaperome differential expression clustering (dendrogram) in B. Turquoise box highlights the human chaperome broken down into functional families in B. Yellow borders indicate marked clusters of chaperome functional family expression and separation of clusters I and II as separated by Euclidean distance clustering of TCGA cancer groups. TCGA cancer group acronyms: THYM (thymoma), ESCA (esophageal carcinoma), BRCA (breast invasive carcinoma), LUAD (lung adenocarcinoma), LUSC (lung squamous cell carcinoma), KICH (kidney chromophobe), STAD (stomach adenocarcinoma), CHOL (cholangiocarcinoma), LIHC (liver hepatocellular carcinoma), PRAD (prostate adenocarcinoma), HNSC (head and neck squamous cell carcinoma), KIRP (kidney renal papillary cell carcinoma), SARC (sarcoma), UCEC (uterine corpus endometrial carcinoma), BLCA (bladder urothelial carcinoma), PAAD (pancreatic adenocarcinoma), CESC (cervical squamous cell carcinoma and endocervical adenocarcinoma), GBM (glioblastoma multiforme), KIRC (kidney renal clear cell carcinoma), SKCM (skin cutaneous melanoma), PCPG (pheochromocytoma and paraganglioma), THCA (thyroid carcinoma).

Besides marked differences in the pattern of cancer functional family changes, Euclidean clustering of cancer groups (rows) revealed two major clusters (Fig 2B). The vast majority of cancers is characterised by the consistent upregulation of HSP90s, ER-specific chaperones, HSP60s, PFDs, HSP100s and MITOs, opposed by a very consistent downregulation of sHSPs and a more cancer-specific overall downregulation of the HSP70-HSP40 system an TPR-domain co-chaperones. This group comprises Cluster I, representing ∼91% of cancers, while Cluster II comprises ∼9% of cancers with largely opposed chaperome deregulation signatures, in this set namely skin cutaneous melanoma (SKCM), and pheochromocytoma and paraganglioma (PCPG).

In summary, systematically assessing gene expression data derived from a total of 10,456 patient samples uncovers broad differences in chaperome-scale deregulation across the variety of human solid cancers. While the vast majority of cancers shows consistent and strong upregulation of chaperome genes, this analysis reveals marked clusters of chaperome functional family expression signatures that further stratify cancers by differential chaperome expression.

### Preferential Upregulation of ATP-Dependent Chaperones in Cancers

The major ATP-dependent chaperone functional families are consistently upregulated across a majority of cancers, while co-chaperones and sHSPs are consistently repressed (Fig 2B). In order to quantify this trend, we assessed the 22 differentially regulated cancer chaperomes for functional characteristics.

First, we compared expression of 88 chaperones against 244 co-chaperones represented in the human chaperome (Brehme et al. 2014). Projecting TCGA cancer groups by their chaperone and co-chaperone differential expression highlights a significant preponderance of cancer chaperome upregulation, including both chaperones and co-chaperones, while only a minor fraction of each is downregulated (S2 Figs A,B). Overall, chaperones tend to be more upregulated than co-chaperones (S2 Figure B). Consistently, within the group of chaperones, we find an overall preponderance of upregulation of both ATP-dependent and ATP-independent chaperones, while only small fractions each are downregulated (S2 Figs C,D). The 50 ATP-dependent chaperones are more upregulated than the 38 ATP-independent chaperones, while ATP-independent chaperones are more downregulated than ATP-dependent chaperones (S2 Figure D).

This analysis exposes a sub-group of cancers as notable exceptions to these trends, suggesting fundamental differences in chaperome deregulation. Projection of TCGA cancer groups by chaperone and co-chaperone up- and downregulation lends support for two groups of cancers, Group 1 and Group 2 (Fig 3A). These groups are recapitulated when projecting cancers by up- and downregulation of ATP-dependent versus ATP-independent chaperones (Fig 3C). K-means clustering confirms the significant separation of the Group 2 cancers pheochromocytoma and paraganglioma (PCPG), thyroid carcinoma (THCA), and the three kidney cancers kidney chromophobe (KICH), kidney renal papillary cell carcinoma (KIRP), and kidney renal clear cell carcinoma (KIRC) from Group 1 cancers, with a median silhouette width of *s* = 0.63 (Fig 3A) and *s* = 0.68 (Fig 3C). Group 1 cancers (red) represent the majority of cancers, characterized by strong overall chaperome upregulation, with low chaperone and co-chaperone repression, and a trend for upregulation of ATP-dependent chaperones (Fig 3B,D). Five Group 2 cancers however partition more distantly, with a lack of chaperome upregulation (Fig 3A,C). These cancers include three different kidney cancers, KICH, KIRP, and KIRC, which consistently lack chaperome upregulation (Fig 3A,B). Also, ATP-dependent chaperones are not preferentially upregulated in kidney cancers. Rather, an inverse trend is observed with increased downregulation of ATP-dependent chaperones (Fig 3C,D). Notably also, pheochromocytoma and paraganglioma (PCPG), rare related tumors of orthosympathetic origin, similarly show an even more prominent inverse alteration, with a preponderance of overall chaperome downregulation and preferential downregulation of ATP-dependent chaperones. Pheochromocytomas originate in the adrenal medulla, with close spatial association to the kidney, whose cancers are also Group 2 cancers. THCA is similar to PCPG, with a preferential downregulation of the chaperome and both PCPG and THCA represent tumors forming from cells of neuroendocrine origin.

**Fig 3.**
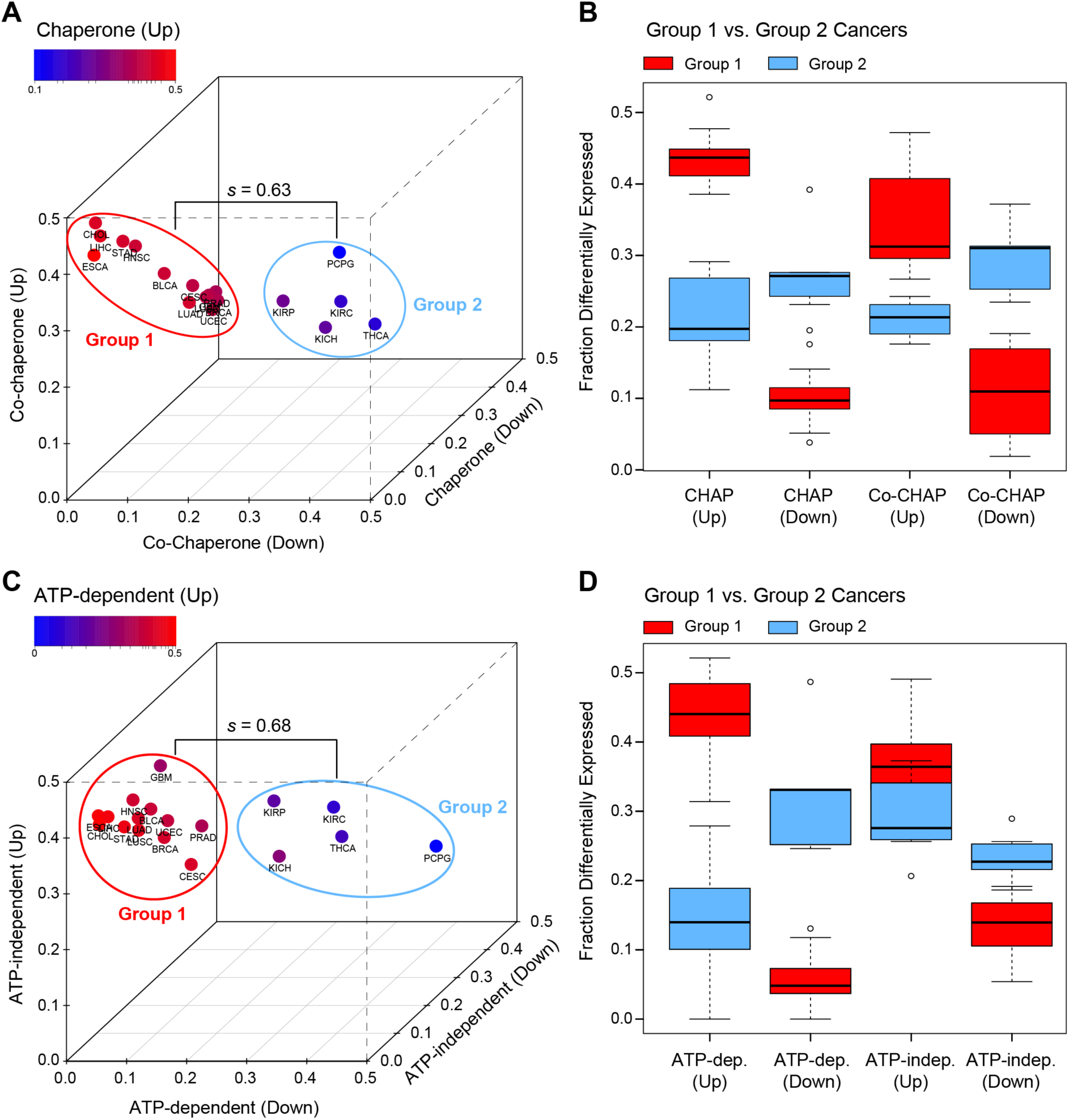
Preferential Upregulation of ATP-Dependent Chaperones in Group 1 Cancers. Analysis of differential cancer gene expression of chaperome functional subsets. **A.** Comparing upregulation and downregulation of gene expression (∆GSA) of chaperones (n = 88) and co-chaperones (n = 244) reveals general chaperome upregulation for the majority of cancers (Group 1), while a small group of Group 2 cancers do not follow this trend. Colour code indicates chaperone up-regulation of gene expression, axes represent chaperone downregulation, co-chaperone upregulation and down-regulation of gene expression. *s =* median silhouette width (k-means clustering). **B.** Box-and-whisker plots highlight fractions of differentially expressed genes in each chaperome subset for Group 1 (red) and Group 2 (blue) cancers separately (see A). Differentially expressed genes in each set were obtained by linear modelling (Limma package in R), considering genes with p value < 0.05 (Benjamini-Hochberg corrected). Box boundaries, 25% and 75% quartiles; middle horizontal line, median; whiskers, quartile boundaries for values beyond 1.5 times the interquartile range; small circle, outlier. **C.** Assessing differential expression of ATP-dependent (n = 50) vs. ATP-independent (n = 38) chaperones highlights preferential upregulation of ATP-dependent chaperones in Group 1 cancers. Group 2 cancers do not follow this trend. *s =* median silhouette width (k-means clustering). **D.** Box- and-whisker plots (as in B.) show fractions of differentially expressed genes in the two sets of ATP-dependent and ATP-independent chaperones, partitioned by Group 1 and Group 2 cancers (see C).

Collectively, the data point to a preferential upregulation of ATP-dependent chaperones in the majority of cancers, which we refer to as Group 1 cancers, with general differences in chaperome deregulation in Group 2 cancers, comprising kidney cancers and cancers of neuroendocrine origin, such as PCPG and THCA.

### Proteasome and TRiC/CCT Upregulation in Cancers Reminisces Enhanced Stem Cell Proteostasis

Human embryonic stem cells (hESCs) are characterized by their capacity to replicate infinitely in culture, while maintaining a pluripotent state (Miura et al. 2004). This immortal, undifferentiated phenotype resembles hallmark features of cancer cells such as an elevated global translational rate (You et al. 2015) and is expected to demand increased PN capacity capable of buffering imbalances to maintain proteostasis. Given the “stemness” phenotype of cancer cells and their resemblances with pluripotent stem cells we hypothesized that the consistent chaperome upregulation in cancers acts to mimic an enhanced stem cell PN setup. Increased proteasome activity (Vilchez et al. 2012) and elevated overall levels of the TRiC/CCT complex (Noormohammadi et al. 2016), representatives of the clearance and folding functional arms of the PN, respectively, have recently been associated with the intrinsic PN of pluripotent stem cells that acts to support their identity and immortality. It can be hypothesized that increased levels of central PN processes in stem cells exemplify characteristics of an enhanced PN setup. We thus assessed to which extent this stem cell PN setup is recapitulated in cancers.

First, we assessed differential changes of the proteasome across TCGA cancers and observed an overall consistent upregulation of the 43 proteasomal genes (HGNC Family ID 690) in > 70% of cancers (Fig 4A, S1 Table B) (Gray et al. 2015), matching the role of increased proteasomal activity for proteome maintenance in stem cells (Vilchez et al. 2012). Notably, Group 1 and Group 2 cancers, which are specifically defined based on chaperome differential expression signatures (Fig 3), do not co-partition with cancer clusters obtained by proteasome differential expression (Fig 4A). Next, we assessed cancer differential expression of the eukaryotic chaperonin TRiC/CCT, a hetero-oligomeric complex of two stacked rings with each eight paralogous subunits representing the cytoplasmic ATP-driven HSP60 chaperones in charge of folding approximately 10% of the proteome (Lopez et al. 2015). TRiC/CCT is highly conserved and essential for cell viability (Lopez et al. 2015). Loss of complex subunits induces cell death and a decline of pluripotency of hESCs and induced pluripotent stem cells (iPSCs) (Noormohammadi et al. 2016). Within the PN, TRiC/CCT mediated folding and autophagic clearance act in concert to prevent aggregation (Pavel et al. 2016). TRiC/CCT levels decline during stem cell differentiation, and CCT8 acts as complex assembly factor (Noormohammadi et al. 2016). Intrigued by the finding that CCT8 is the most highly elevated subunit in stem cells and likely acting as assembly factor (Noormohammadi et al. 2016), we assessed differential expression of individual TRiC/CCT subunits across cancers. Hierarchical clustering of subunit expression across solid cancers highlighted CCT8 as highly consistently upregulated across all cancers (mean change = 0.76, *t* test), and as overall second most highly upregulated subunit besides CCT6A (mean change = 0.786, *t* test) (Fig 4B). Consistent with the overall preferential downregulation of ATP-dependent chaperones observed in Group 2 cancers (Fig 3), we found these cancers to cluster together with overall lowest TRiC/CCT expression, where PCPG stands out with a consistent downregulation of all subunits (Fig 4B).

**Fig 4.**
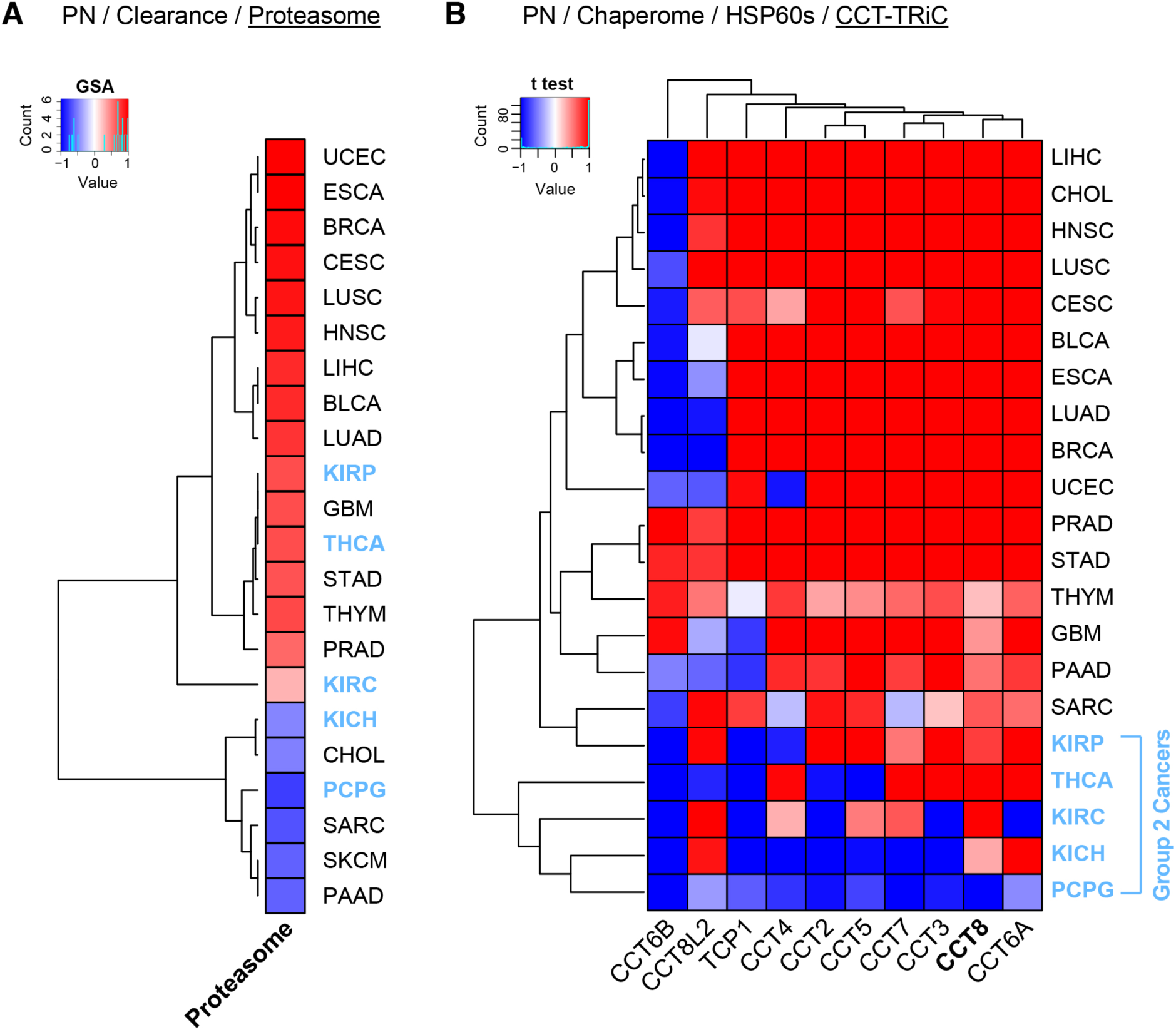
Proteasome and TRiC/CCT Increase in Human Cancers Reminisces Stem Cell Proteostasis. **A.** Heatmap indicates overall gene expression changes (∆GSA) of the human proteasome (43 genes, HGNC Family ID 690) throughout 22 TCGA solid cancers. Heatmap ∆GSA values are in the interval [-1, +1], where ‘+1’ indicates significant upregulation (p value = 0), while ‘-1’ indicates significant downregulation (p value = 0) as in Fig 2. **B.** Heatmap highlights HSP60 gene level differential expression of TRiC/CCT complex subunits throughout 22 TCGA solid cancers. Heatmap indicates significance of up- or downregulation of gene expression (*t* test) in cancer compared to matching healthy tissue (1 - signed p value) in the interval [-1, +1], where ‘+1’ indicates significant upregulation (p value = 0), while ‘-1’ indicates significant downregulation (p value = 0). Blue highlights indicate Group 2 cancers (KICH, KIRC, KIRP, PCPG, and THCA).

Together these findings suggest that proteostasis shifts in cancer cells add to an altered, enhanced PN state that mimics the immortal and resilient stem cell phenotype, buffering genome instability and ensuing proteomic imbalances in support of sustained and increased cellular proliferation throughout cancerogenesis.

### Opposing Chaperome Deregulation in Cancers and Neurodegenerative Diseases

Overall chaperome upregulation in cancers, with preferential enrichment for upregulation of ATP-dependent chaperones, alongside consistent downregulation of sHSPs, is diametrically opposed to chaperome deregulation trends previously observed in a study of chaperome alterations in human aging brains and in patient brains with age-onset neurodegenerative diseases (Brehme et al. 2014). While sHSPs were the only chaperome family found significantly induced in brain aging and the age-onset neurodegenerative diseases Alzheimer’s (AD), Huntington’s (HD), and Parkinson’s (PD) disease, this family is consistently downregulated across cancers (Fig 2B). This opposed chaperome deregulation points towards characteristic and fundamental differences in PN deregulation between disease families.

To investigate this disease group difference further, we applied the analysis outlined for cancers above also on the gene expression datasets that had earlier revealed global chaperome repression in AD, HD, and PD (Brehme et al. 2014). Our analysis reproduced the human chaperome as overall downregulated across AD, HD, and PD as compared to random permutations of non-chaperome genes (-0,20 ∆GSA group mean change, Fig 5A). Delving deeper into chaperome functional subfamilies, we reproduce earlier findings reporting broad repression of the major chaperome functional families except for sHSPs, the only family found strongly upregulated (+0,76) accompanied by slight upregulation of ER-specific chaperones (+0,21) and TPRs (+0,13) (Fig 5B). Thereby, our analytical workflow reproduces previously observed trends obtained in independent analyses, with different methods. With strong sHSP repression and upregulation of the HSP90, ER, HSP60, PFD, HSP100 and MITO chaperone families, cancers and neurodegenerative diseases display markedly diametrically opposed chaperome deregulation, not only at the overall chaperome-level (Fig 5C), but also with respect to alteration trends of chaperome functional families, where 70% of functional groups are altered in opposite directions (Fig 5D).

**Fig 5.**
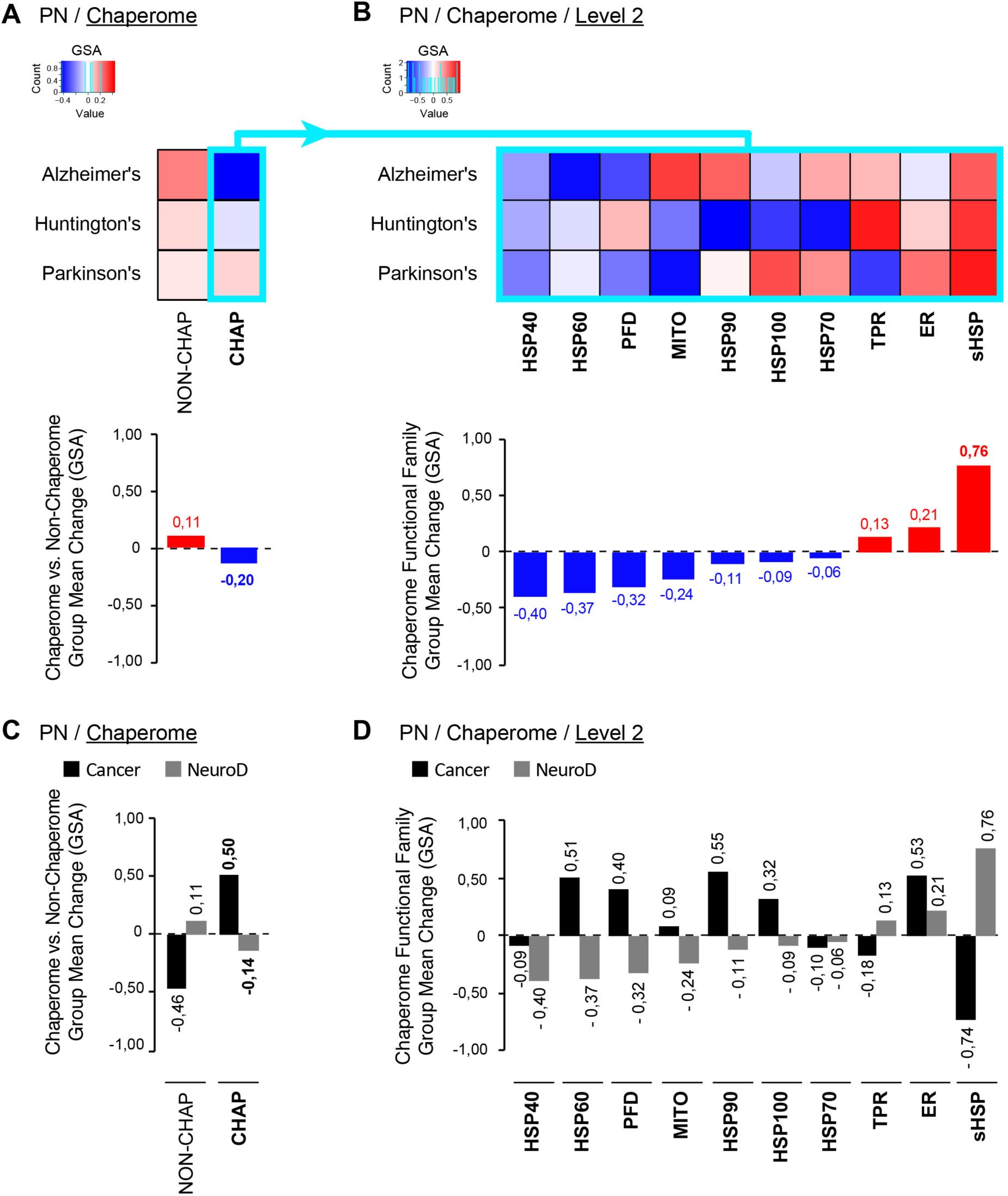
Opposing Chaperome Deregulation in Cancers and Neurodegenerative Diseases. **A - B.** Heatmaps indicate significance of up or down-regulation of gene expression (∆GSA) of chaperome vs. non-chaperome genes (**A**) and chaperome functional families (**B**) in Alzheimer’s (AD), Huntington’s (HD), and Parkinson’s disease (PD) compared to age-matched healthy controls. Datasets: GSE5281 (AD, superior frontal gyrus) (Liang et al. 2008), GSE3790 (HD, nucleus caudatus) (Hodges et al. 2006), and GSE20295 (PD, substantia nigra) (Moran et al. 2006). ∆GSA values are in the interval [-1, +1], where ‘+1’ indicates significant upregulation (p value = 0), while ‘-1’ indicates significant downregulation (p value = 0) as in Fig 2. Functional families (columns) are ordered by increasing ∆GSA group mean change (bar graphs). Turquoise box highlights the human chaperome. **C - D.** Side-by-side comparison of gene expression changes (∆GSA) of chaperome vs. non-chaperome (C), and chaperome functional families (D) in cancers versus neurodegenerative diseases (NeuroD). Bar graphs show ∆GSA group mean changes in cancers (black) compared to NeuroD (grey).

These opposing chaperome deregulation signatures are in line with differing implications of proteostasis alterations in these diseases. While broad chaperome repression and proteostasis functional collapse is associated with aggregation and cytotoxicity of chronically expressed misfolding-prone proteins in neurodegenerative diseases (Brehme et al. 2014), enhanced proteostasis buffering capacity is associated with “stemness”, immortality and proliferative potential of both stem and cancer cells (Whitesell and Lindquist 2005). Indeed, epidemiological evidence suggests an inverse correlation between cancers and neurodegenerative diseases (Driver et al. 2012, Musicco et al. 2013, Ou et al. 2013, Li et al. 2014), supportive of a potential mechanistic link between opposed chaperome deregulation and the molecular underpinnings of the two disease groups. These global differences in chaperome deregulation call for a systematic and quantitative assessment of PN deregulation dynamics in human diseases.

### A Multi-Step Dimension-Reduction Approach Enables Quantitative Visualisation of Chaperome Shifts in Cancers

In light of the diverse signatures of differential chaperome deregulation observed across cancers (Fig 2), and motivated by the increasing amount of genomics datasets available for cancers and other human diseases, we aimed at reducing data complexity by extracting quantitative indicators of chaperome differential cancer gene expression alterations, in order to gain insights through reduced complexity while retaining maximum information content.

We devised Meta-PCA, a novel principal component analysis (PCA) based semi-supervised two-step dimension reduction approach that facilitates stratification of cancer patient and normal control samples within heterogeneous gene expression datasets (Fig 1B). Based on previous work on dimensionality reduction of heterogeneous gene expression datasets (Lenz et al. 2013, Lenz et al. 2016), we hypothesized that the underlying information contained in each chaperome functional group has a low dimensionality and could be surfaced using PCA, if there were sufficient samples available to represent the complete heterogeneity of chaperome alterations in cancers. Compared to conventional PCA, our method can deal with the effect of different group sizes, which as a confounder would negatively affect PCA results. Compared to simply calculating mean expression values of each functional group’s genes, Meta-PCA considers a wider range of gene expression information inherent to each functional group, resulting in scores with higher resolution. This could also be achieved by fully supervised methods such as Linear Discriminant Analysis. However, this case requires use of a method, which is blind to sample annotations so that these can later be used for validation, such as the unsupervised classification of cancer tissue type. Meta-PCA first uses tissue-wise PCA analyses to separate cancerous from control samples for individual tissues and maps functional family group gene expression changes to the global, or “meta”, mean expression change across cancers in order to obtain M-scores as quantitative indices of relative disease gene expression change (Equation 1). Inherent to the Meta-PCA method, patient-specific genomic variability is averaged out through the use of Meta-PCs derived from the TCGA collection of cancer biopsy samples, yielding mean reference boundaries. Assessing quantitative M-scores obtained through Meta-PCA against differential gene expression (∆GSA) obtained via GSA (Fig 2), we observed an overall significant correlation of 0.61 (Limma package) (S1 Figure). In addition to general comparability between results obtained through both methods, Meta-PCA analysis reduces complexity while retaining genomic information. Therefore, we focus on Meta-PCA for quantitative representation of proteostasis alterations in human diseases. We plot differential chaperome changes as M-scores for all cancer samples and chaperome functional families simultaneously using polar plots, such that axes represent functional families or sub-groups (Fig 2B). We obtain the mean of all biopsy samples as reference boundaries for healthy (blue line) and cancer groups (red line) and include the 90% confidence interval (CI) (red and blue halos) (Fig 6). This quantitative visualisation reduces complexity and highlights relativity of disease gene expression changes at a chaperome-scale. The polar maps recapitulate characteristic chaperome deregulation signatures in GSA-derived clusters of functional family upregulation and downregulation signatures, for instance in Cluster I cancers such as lung adenocarcinoma (LUAD) (Figs 2, 6A), or the inverse trend with overall chaperome downregulation in Cluster II cancers such as pheochromocytoma and paraganglioma (PCPG) (Figs 2, 6B). Concordantly, chaperome polar maps reveal characteristic patterns of cancer groups stratified based on differential expression of ATP-dependent chaperones versus ATP-independent co-chaperones, Group 1 versus Group 2 cancers (Figs 3, 6). In LUAD, representative of Group 1 cancers, most functional chaperome families, with a preferential enrichment of ATP-dependent chaperones, are upregulated, while sHSPs are reduced (Fig 6A). On the contrary, in Group 2 cancers such as PCPG, gene expression of most chaperome functional families is downregulated (Fig 6B). Inconsistencies between these broad clusters exist, suggesting differences in tissue of origin and molecular underpinnings of respective cancers. However, broad commonalities between distinct cancers originating from the same organ are revealed. For instance, lung adenocarcinoma (LUAD) and lung squamous cell carcinoma (LUSC) share overall similarity, revealing only subtle differences, for instance in HSP40 expression (Figs 2, S3). The kidney cancers KICH, KIRP, and KIRC also show similar patterns. As Group 2 cancers, they share and stand out against other cancers with a lack of preferential upregulation of ATP-dependent chaperones (Fig 3), and overall reduced upregulation, or downregulation, of HSP60s (Figs 2, 3, 4B, S3). A recent study indeed implicated HSP60 downregulation in tumorigenesis and progression of clear cell renal cell carcinoma (KIRC) by disrupting mitochondrial proteostasis (Tang et al. 2016).

**Fig 6.**
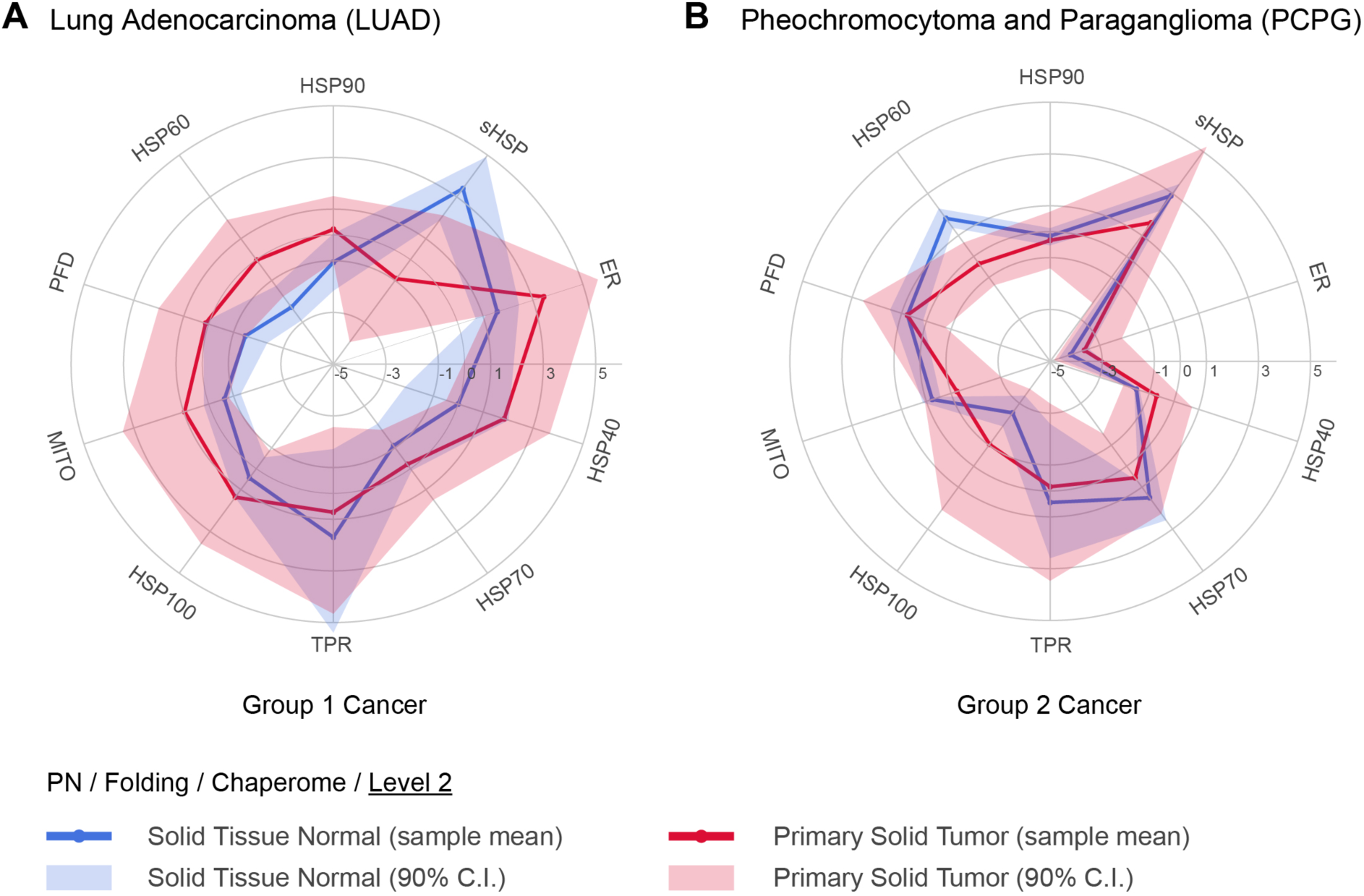
Polar Maps of Chaperome Shifts in Human Cancers. Polar plot visualization as novel quantitative and contextual representation of M-scores, new Meta-PCA derived quantitative indices of relative disease gene expression shifts of chaperome functional processes compared to normal tissue counterparts. Examples representative of Group 1 cancers, lung adenocarcinoma (LUAD) (**A**), and Group 2 cancers, pheochromocytoma and paraganglioma (PCPG) (**B**) are shown. Blue (normal) and red lines (cancer), sample means. Halos, confidence interval at the 90% quantile range (5% - 95%). LUAD (lung adenocarcinoma), PCPG (pheochromocytoma and paraganglioma).

Overall, these contextual quantitative representations enable an appreciation of the complex chaperome shifts in different cancer tissues derived from > 10,000 patient biopsy samples. The resulting compendium of differential cancer chaperome polar plots (S3 Figure) is also available online through the Proteostasis Profiler (Pro^2^) tool associated to this study.

### Interactome-guided Topographic Maps Highlight Relative Changes of Chaperome Functional Families in Cancers

The integration of disease-related differential transcriptomic changes with the cellular protein interactome network, or the edgotype, is instrumental to our understanding of genotype - phenotype relationships (Sahni et al. 2013). Towards integrated quantitative views of cancer chaperome deregulation, we curated a high-confidence physical chaperome protein-protein interactome network (CHAP-PPI) to serve as coordinate base grid layout for the analysis of differential chaperome topographies (Fig 1C).

We started with 328,244 unique human PPIs (edges) between 16,995 proteins (nodes) downloaded from the BioGRID, IntAct, DIP, and MINT databases (Chatr-Aryamontri et al. 2013),(Kerrien et al. 2012),(Salwinski et al. 2004),(Licata et al. 2012). Zooming in on cancer chaperome alterations in the context of physical interactome wiring, we extracted the CHAP-PPI considering the 332 human chaperome genes as previously described (Brehme et al. 2014). Considering edges with the PSI-MI annotation ‘physical association’ we obtained 272,367 unique physical edges, of which 666 unique edges connect 220 chaperome nodes. We developed a custom script to curate the high-confidence physical CHAP-PPI, considering edges with multiple pieces of evidence, either experimental methods or publications (PMIDs), as more reliable than those supported by only a single piece of evidence. The curation script resolves ambiguous database annotation of methods terms through up-propagation within the PSI-MI ontology tree, only accepting uniquely different or rejecting identical experimental evidence. Automated interactome curation results in eight curation levels (L1 - L8), through which we obtain three interactomes of increasing confidence level (see Methods). All 666 unique physical edges between 220 chaperome nodes, without curation for type or number of evidence, represent the single evidence chaperome interactome (SE-CHAP). Curating for high-confidence interactions, we obtained a multiple evidence chaperome (ME-CHAP) comprised of 222 unique physical chaperome edges between 128 chaperome nodes, of which a subset of 132 interactions between 96 nodes is supported by multiple different experimental methods (MM-CHAP) (S2 Table).

In order to enable focussed views on transcriptomic alterations of top-level chaperome functional families, we collapsed individual nodes onto functional family meta-nodes, and edges shared between families were collapsed as meta-edges such that meta-node sizes correspond to the number of family members and meta-edge thickness represents the number of shared interactions between families. We considered the meta-interactome derived from the curated high-confidence ME-CHAP interactome (S2 Table), where all meta-nodes corresponding to the 10 functional chaperome families are fully inter-connected in a single network component. We set node colour to visualize cancer gene expression changes (M-scores) and applied a force-directed spring layout algorithm to optimize graph layout (Kamada and Kawai 1989). The resulting integrated cancer chaperome meta-interactomes visualize relative chaperome differential changes at reduced complexity across diverse human cancers in the context of physical interactome connectivity (Figs 7A, S4). Next, we extract x-y coordinates of the chaperome meta-nodes in the optimized meta-network graph to serve as a 2-dimensional base grid (x-y coordinates) guiding the spatial layout of 3-dimensional chaperome topographic maps of differential chaperome gene expression changes (M-scores) between cancerous and healthy biopsies (z coordinate) (Figs 7B, S5).

**Fig 7.**
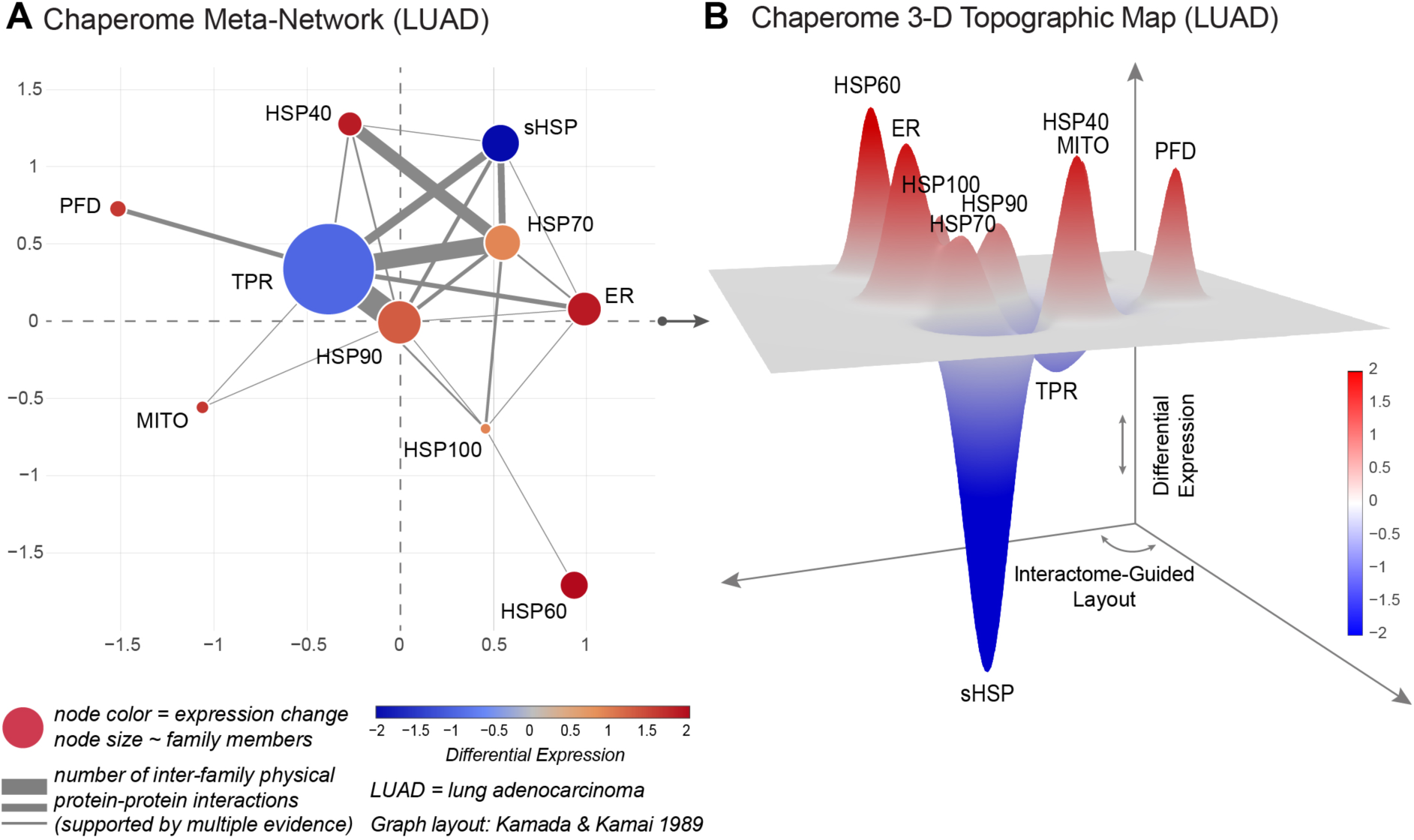
Interactome-Guided Topographic Maps of Cancer Chaperome Shifts. **A.** Cancer-specific chaperome meta-interactome networks, collapsing network nodes and edges onto meta-nodes and meta-edges, highlight cancer-specific gene expression changes of chaperome functional families in context of connectivity within the high-confidence physical interactome network (ME-CHAP) at reduced complexity. Node size and edge thickness correspond to the number of functional family member nodes and the sum of inter-family edges, respectively. Node colour indicates combined cancer gene expression changes based on Meta-PCA derived M-scores. LUAD = lung adenocarcinoma. **B.** Projecting differential changes between cancer and normal counterpart biopsy gene expression based on Meta-PCA derived M-scores (z dimension) onto the ME-CHAP interactome derived meta-network (see A) serving as base-grid layout (x-y dimensions), we derive cancer-specific interactome-guided 3D topographic maps. LUAD = lung adenocarcinoma. Both visualisation, meta-networks (A) and 3D-topographic maps (B) are accessible through Proteostasis Profiler (Pro^2^).

This interactome-guided topographic display of differential chaperome alterations enables dimensionality and complexity reduction for the coherent display and comparative analysis of functional network shifts that can serve to compare differential changes i) in disease versus controls, ii) between diseases and disease classes, and iii) between perturbed or unperturbed states across large numbers of heterogeneous genomic datasets. Furthermore, this visualization lends itself for a systems-level assessment of PN deregulation topologies and their readjustment in human disease and therapeutic intervention. We implemented topographic map visualisations into the Proteostasis Profiler (Pro^2^) suite of tools, to improve accessibility and applicability by the scientific community.

### Proteostasis Profiler (Pro^2^) - An Integrated Online Resource and Toolbox for the Analysis and Visualization of Proteostasis Disease Alterations

Here we exemplify a systematic analysis of differential chaperome gene expression alterations in cancers and neurodegenerative diseases. We reduce complexity through the focus on top-level chaperome functional families. The challenge in this analysis is in the complexity and heterogeneity of available samples for disease groups such as cancers, combined with the multitude of diverse biological processes interconnected within the PN and within its functional processes, as highlighted here at hands of the human chaperome.

To date, there has been no systematic interactome-guided analysis of the implications and alterations of cellular proteostasis biology at a systems-level, in a comprehensive set of diseases, such as cancers. Here, we showcase an integrated analytical workflow for the dimension reduction, analysis and visualization of chaperome differential alterations in a representative set of human solid cancers. Our approach focuses on the visualisation of a confined set of Meta-PCA derived quantitative M-scores as descriptors of top-level chaperome functional families. We have developed “Proteostasis Profiler” (Pro^2^) as an integrated web-based resource and suite of tools, for interactive dimensionality-reduction, analysis and visualisation of disease-specific alterations of proteostasis functional arms, such as the chaperome, in the context of the interactome network. In this study we highlight Pro^2^ use-cases for the human chaperome across TCGA solid cancers in comparison to the three major neurodegenerative diseases using differential gene expression heat maps (∆GSA) (Figs 2, 5), Meta-PCA derived quantitative polar plots (M-scores) (Figs 6, S3), meta-interactomes and interactome-guided 3-D topographic maps (Figs 7, S4, S5). Pro^2^ provides an integrated online suite for the application of the underlying algorithms. Pro^2^ is accessible directly at http://www.proteostasys.org or through the JRC-COMBINE resources collection at http://www.combine.rwth-aachen.de/index.php/resources.html.

## Discussion

Cancer prevalence, genetic complexity and heterogeneity represent unmet medical need and a significant challenge to personalized medicine, calling for genome-informed therapeutic intervention strategies (Burrell et al. 2013). While important progress has been made in the elucidation of proteostasis alterations in human diseases, revealing numerous alterations of PN functional processes not only in neurodegenerative or metabolic diseases but also in cancers, paradoxically the characteristics and extent of PN alterations in cancers are largely unexplored and not understood at a systems-level. Cancer cell line global transcriptional characteristics have been extensively studied (Klijn et al. 2015) and numerous individual studies have assessed alterations of various chaperone and co-chaperone expression levels in specific cancers (Jolly and Morimoto 2000, Whitesell and Lindquist 2005). In light of limitations in the clinical translation of hypotheses derived from cell lines and the lack of a systems-level understanding of proteostasis alterations in human disease, we argue that precise quantitative maps of proteostasis deregulation in human disease derived directly from clinical biopsy data will enable precise understanding of the role of PN alterations in pathogenesis towards testable hypotheses and rationalised approaches of PR therapy (Balch et al. 2008, Hutt and Balch 2010).

Here, we focused on the human chaperome, a central PN component, and highly conserved facilitator and safeguard of the healthy folded proteome using an expert-curated human chaperome functional gene ontology comprising an ensemble of 332 chaperone and co-chaperone genes (Brehme et al. 2014) to systematically characterize chaperome alterations in a representative clinically relevant dataset of 22 human solid cancers with matching healthy tissue, corresponding to over 10,000 patient biopsy samples provided through the TCGA consortium (Cancer Genome Atlas Research et al. 2013). We found the human chaperome to be consistently highly upregulated across the vast majority of cancers assessed. While numerous individual chaperones and co-chaperones have previously been found upregulated in individual cancers (Jolly and Morimoto 2000, Whitesell and Lindquist 2005), this knowledge has not been coherently derived from consistent data resources or systematic genome-wide analyses in biopsy tissue before. Here, we provide systematic quantitative maps of chaperome deregulation in cancers that highlight the relevance, characteristics and extent of chaperome upregulation in cancers. Our analysis revealed chaperome deregulation signatures that not only feature broad upregulation of ATP-dependent chaperones but also consistent repression of ER-specific chaperones and the ATP-independent sHSPs. The data also suggest two cancer groups that can be stratified specifically by their chaperome deregulation patterns. Overall chaperome upregulation across cancers is in agreement with existing evidence on individual chaperones that has been previously reviewed (Morimoto 1991, Fuller et al. 1994, Jolly and Morimoto 2000). For instance, elevated heat shock protein expression levels have been reported for HSP90 in breast and lung cancers (Jameel et al. 1992, Wong and Wispe 1997), HSP70 was found increased in breast, oral, cervical and renal cancers (Ciocca et al. 1993, Kaur and Ralhan 1995, Ralhan and Kaur 1995, Santarosa et al. 1997), and HSP60 showed increased expression in Hodgkin’s disease (Hsu and Hsu 1998). The cellular safeguarding functions of chaperones are subverted during oncogenesis to facilitate malignant transformation in light of increased translational flux and aberrant protein species in cancer cells (Whitesell and Lindquist 2005). Increased chaperone levels have previously been correlated with poor prognosis and cancer survival (Jameel et al. 1992, Yano et al. 1996, Jolly and Morimoto 2000). Chronic dependency on stress response and quality control mechanisms drives cancer cells into a phenotype of non-oncogene addiction (Solimini et al. 2007). The observed extent of chaperome alterations suggests a broader state of cancer “chaperome addiction”, beyond the dependency on individual chaperones.

Evidence points towards functional associations between increased proteostasis buffering capacity and maintenance of “stemness”, immortality and proliferative potential in both cancer cells and pluripotent stem cells (Whitesell and Lindquist 2005). For instance, autophagy was found to maintain “stemness” by preventing senescence through sustained proteostasis (Garcia-Prat et al. 2016). Increased proteasomal activity and elevated levels of the HSP60 chaperonin complex TRiC/CCT have recently been linked to stem cell identity by conferring proteostasis robustness (Vilchez et al. 2012, Noormohammadi et al. 2016). Fundamental similarities between stem cells and cancer raise the question to the extent of similarity between cancer and stem cell PN states and capacity. Our data suggest that cancers consistently display signatures of elevated proteostasis functional processes such as the chaperome and proteasome-mediated clearance, and are in agreement with the hypothesis that upregulated clearance mechanisms such as the proteasome and increased chaperome topologies, particularly increases in ATP-driven chaperones such as the HSP60 chaperonin complex TRiC/CCT, confer increased proteostasis capacity and survival benefits to cancer cells just like they are essential to stem cell biology. Precise knowledge of systems-level network deregulation therefore sheds light on fundamental processes at play from stem cell biology to cancerogenesis. Chaperone upregulation is largely regulated through heat shock factor 1 (HSF1) (Dai et al. 2007). Overexpression of the TRiC/CCT subunit CCT8 protects against *hsf-1* knockdown in *C. elegans* (Noormohammadi et al. 2016), consistent with a regulatory connection between TRiC/CCT and HSF1 (Neef et al. 2014). Connecting processes at the PN level, this evidence suggests a connection between TRiC/CCT and HSF1 stress response signalling also in cancers (Mendillo et al. 2012, Noormohammadi et al. 2016). While increased expression of TRiC/CCT subunits has been observed in cancer cell lines (Boudiaf-Benmammar et al. 2013), and increases in CCT8 expression are linked to individual cancers (Huang et al. 2014, Qiu et al. 2015), we describe consistent TRiC/CCT upregulation within global cancer chaperome signatures throughout the majority of TCGA solid cancers, or Group 1 cancers, whereas Group 2 cancers lack chaperome and, to large extent, TRiC/CCT upregulation.

Contrary to stem cell proteostasis, which is set up to maintain pluripotency and proliferative capacity, neurodegenerative diseases such as Alzheimer’s (AD), Huntington’s (HD), and Parkinson’s disease (PD) display signs of proteostasis functional collapse. Misfolding diseases feature overexpression of aggregation - prone proteins such as Aβ in AD, α-synuclein in PD, or huntingtin in HD that entail a “toxic-gain-of-function” resulting in chaperome overload, gradually exceeding proteostasis capacity (Knowles et al. 2014), while “loss-of-function” misfolding diseases feature specific perturbations such as dysfunctional ∆F508-CFTR in cystic fibrosis (Chanoux and Rubenstein 2012). Functionally deficient steady-state dynamics of the folding environment affect cellular protein repair capacity and proteome maintenance (Wang et al. 2006). Most cancer cells however harbour manifold genetic aberrations even at the karyotype level that likely entail dramatic effects on proteome balance (Harper and Bennett 2016). The collective damage caused by oncoprotein expression, compromised DNA repair, genomic instability, reactive oxygen species (ROS), elevated global translation and chaperome overload triggers stress response mechanisms in light of a challenged cellular proteostasis capacity (Csermely 2001). Chaperome deregulation dynamics observed in cancers indeed display concordantly opposed trends as compared to alterations in the major neurodegenerative diseases.

A recent study linked repression of ∼30% of the human chaperome in aging brains and in neurodegenerative diseases to proteostasis functional collapse and pointed to the role of a chaperome sub-network as a conserved proteostasis safeguard (Brehme et al. 2014). Intriguingly, while only ∼8% of the human orthologous chaperome had protective phenotypes upon functional perturbation in *C. elegans* models of amyloid β (Aβ) and polyQ proteotoxicity, chaperones and co-chaperones far less well studied than HSP90 had equally strong protective effects (Brehme et al. 2014). Similarly, an overlap between the chaperome and the “essentialome” set of 1,658 core fitness genes in K562 leukemia cells (Wang et al. 2015) found only 55 overlapping with the 332 chaperome genes (Balchin et al. 2016). Interestingly, HSP60s showed the highest fraction of essential chaperones in agreement with their function as a highly conserved folding complex that hosts ∼10% of the proteome’s clients (Lopez et al. 2015). Collectively, these findings suggest a highly functionally redundant and robust role of the central conserved chaperome within the PN (Brehme and Voisine 2016).

In summary, our study showcases a systematic profiling of the extent of chaperome deregulation, as a central PN functional arm, in a panel of human cancers and three major neurodegenerative disorders, accompanied by a resource of quantitative multi-dimensional maps with reduced complexity. Therapeutic PN regulation for increased or restored proteostasis capacity may be beneficial in both loss-of-function and gain-of-toxic-function diseases of protein misfolding (Powers et al. 2009). Attenuating the PN on the other hand, such as inhibiting chaperones like HSP70 and HSP90 or the UPS clearance machinery, are widely acknowledged as promising therapeutic avenues in cancers (Whitesell and Lindquist 2005, Miyata et al. 2013, Balchin et al. 2016, Li et al. 2016). While this manuscript was in preparation, Rodina and co-workers reported findings on a highly integrated chaperome subnetwork, or ‘epichaperome’, as a classifier of cancers with high sensitivity to HSP90 inhibition, while cancers with a less interconnected chaperome are less vulnerable by HSP90 inhibition (Rodina et al. 2016). Several HSP90 inhibitors have shown encouraging results in clinical trials (Khajapeer and Baskaran 2015). Our study further supports the central role of the chaperome in PN biology, justifying particular focus on understanding chaperome alterations in human diseases at a systems level. The characteristic signatures of cancer chaperome alterations revealed in this study suggest broad commonalities and differences that could serve as testable hypotheses for therapeutic chaperome targeting strategies in cancer. Our results underline the value of charting quantitative systems-level maps and provide a resource towards an improved functional understanding of proteostasis biology in health and disease. A systems-level understanding of contextual PN alterations throughout the human diseasome will be instrumental for charting a clearer picture of the PN as a therapeutic target space, and as a resource for clinical biomarkers, including the chaperome. In face of increasing amounts of genome-scale disease data we are confronted with tremendous challenges of data complexity. Therefore, our study provides Proteostasis Profiler (Pro^2^), an integrated web-based suite of tools enabling processing, analysis and visualisation of proteostasis alterations in human diseases at reduced dimensionality, towards hypotheses-building for mechanistic understanding and clinical translation.

## Materials and Methods

### Gene Expression Data Preparation

Focussing on pan-cancer analysis of the human chaperome, we chose The Cancer Genome Atlas (TCGA) as the main source for our analyses, as an established dataset that is widely used and adopted by the scientific community. The Broad Institute TCGA GDAC Firehose was accessed to download TCGA RNAseqv2 raw counts data followed by application of the voom method for the transformation of count data to normalized log2-counts per million (logCPM) (Law et al. 2014). Each of these logCPM values were centered gene-wise for sample normalization and comparability and used for all analyses. Considering TCGA clinical data annotation, we extracted those 22 tissue biopsy group datasets that provide both “primary solid tumor” and “solid tissue normal” sample type annotations.

### Gene Set Analysis (GSA) and Heatmap Representation

We applied Gene Set Analysis (GSA) (Efron and Tibshirani 2007), an advanced derivative of Gene Set Enrichment Analysis (GSEA) (Subramanian et al. 2005), in order to assess chaperome gene family expression changes between cancerous and corresponding healthy tissue samples. When applying GSA, we implemented 100 permutations of chaperome genes contained in each functional family in order to allow for statistical assessment of differential expression upon re-standardization of gene groups for more accurate inferences. When applying GSA to the chaperome as one set in comparison to the whole genome (non-chap set), we randomly sampled 332 genes from the whole genome, excluding chaperome genes, and compared them to the 332 chaperome genes in order to exclude bias on group sizes in the comparisons. We applied this random sampling process 100 times in addition to 100 permutations we had on each GSA calculation. We calculated the mean value of all results as a robust measure of chaperome changes with respect to the genome. Results are displayed as heatmaps indicating significance of up or down-regulation of gene expression as ∆GSA values derived from the difference of (1 - upregulation p value) - (1 - downregulation p value) in disease compared to matching healthy tissue for TCGA cancer datasets, or control patient biopsies for neurodegenerative disease datasets (AD, HD, PD). ∆GSA values are normalized within the interval [-1, +1], where ‘+1’ indicates significant upregulation (upregulation p value = 0), while ‘-1’ indicates significant down-regulation (downregulation p value = 0), accordingly. Bar graphs represent group mean changes of each chaperome functional family gene group over all diseases.

### Linear Modeling of Chaperome Functional Subsets

We subdivided the human chaperome into functional subsets of chaperones and co-chaperones, and further divided chaperones into two sets of ATP-dependent and ATP-independent chaperones according to the annotations provided by Brehme et al. 2014 (Brehme et al. 2014). We performed linear modelling using the Limma package in R. Genes with p values < 0.05 following Benjamini-Hochberg correction are considered in the fraction of differentially expressed genes corresponding to each functional subset.

### Meta-Principal Component Analysis (Meta-PCA)

Gene Set Analysis (GSA) is a statistical hypothesis testing method that is by definition limited to confirmatory data analysis with respect to pre-existing hypotheses. In order to serve the goal of quantitative exploratory pan-cancer chaperome analysis, while retaining a maximum information content during model reduction, we devised Meta-PCA, a novel quantitative multi-step dimension reduction model fitting strategy based on principal component analysis (PCA). Principal component analysis (PCA) uses orthogonal transformation to convert a set of variables to linearly uncorrelated variables, such that they are ordered by their information content, which allows for removal of dimensions with lowest information content for dimensionality reduction in complex heterogeneous datasets. In order to stratify cancer from healthy biopsy gene expression samples based on chaperome functional family gene expression in highly convoluted datasets comprising multiple different cancer types, we designed Meta-PCA as a novel two-step method capable of handling this type of heterogeneous data. We hypothesized that each chaperome functional family or process can be described by a low number of variable dimensions, considering that genes within each group are either related or act together in molecular complexes. Therefore, we used a PCA-based approach for quantitative assessment and dimensionality reduction of functional chaperome alterations based on disease gene expression data. Challenged by highly varying sample counts in the different TCGA cancer group datasets, where datasets (tissues) with high sample numbers are at risk of dominating PCA results as compared to cancer groups with low sample numbers, we developed a custom approach that is not limited by a lack of underlying models for interpolation or undesirable loss of information, such as in up- or down-sampling, respectively, allowing us to consider all samples in the included TCGA cancer groups. Assuming distinct roles for each chaperome functional group we define

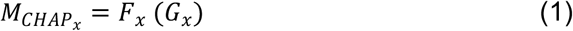

where *M* denotes the M-score of chaperome (CHAP) family *x*, *G*_*x*_ is the vector of gene expression values corresponding to genes in CHAP family *x*, and *F*_*x*_ is the function we want to fit. For simplicity, we considered a linear first degree model as follows: *F*_*x*_ is a vector of weights *W*_*x*_ with identical length as the vector *G*_*x*_, and we aim to find *W*_*x*_ for all *x* using PCA. Assuming equivalent biological function of each *CHAP*_*x*_ among all tissues, we first calculate *F*_*x*_ for each tissue in order to separate disease from healthy samples for each tissue, and then combine all “relevant” PCs in order to obtain the main underlying PC, or ‘Meta-PC’, of the corresponding CHAP group. We outline the ‘Meta-PCA’ algorithm as follows:

**Step 0**: For each CHAP group and tissue we assume a model

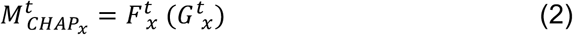

Where 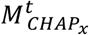 is the M-score of CHAP group x in tissue t, 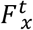 is the unknown function mapping gene expression values for CHAP group *x* in tissue *t* to an M-score value, and 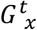 is the gene expression vector of all genes in CHAP group *x* in tissue *t*.

**Step 1:** Assuming 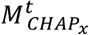 can be approximated using PC1, we assume 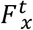 is equal to 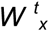 which is the vector of weights for CHAP group *x* and tissue *t*. Then we calculate PCA on the gene expression matrix (GEX) comprising all genes in CHAP group x, and all samples of tissue *t*, including ‘solid tissue normal’ and ‘primary solid tumors’. So in this step we have *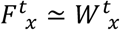* as loadings of PC1.

**Step 2**: The 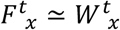 assumption in Step 1 is not necessarily true; PCA extracts the most variable direction in GEX, but in case CHAP group *x* does not change drastically between healthy and cancer, PC1 will represent an unwanted variable or even noise. So we have to filter out the *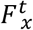* that did not fit well to the data. For this we use Student’s *t*-test. For each tissue, we test the separation of 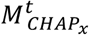 between ‘solid tissue normal’ and ‘primary solid tumor’ samples, and discard all 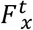 with p values > 10^−4^.

**Step 3**: We combine all 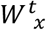 to obtain *W*_*x*_, which is the universal mapping of gene expressions in CHAP group *x* to its corresponding M-score, regardless of tissue type. Therefore, we calculate *W*_*x*_ as

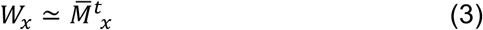

where the loading of each gene in the universal mapping is the mean value of all the loadings of the same gene on different tissues. Importantly, prior to calculating mean loadings, we set all 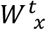 to be uni-directed in order preserve directionality of change from healthy to cancer, yielding final Meta-PCs. *W*_*x*_ can be used as the universal function *F*_*x*_ (Equation 1) in order to map a query sample to the corresponding M-score of CHAP group *x*.

**Step 3’**: In order to validate *F*_*x*_ and resulting M-scores we performed random forest regression using 80% of M-scores and their annotation labels as training set and 20% as test set.

### Quantitative Visualisation of Chaperome Alterations in Diverse Cancers

In order to visually represent quantifications of chaperome functional family differential cancer gene expression, we used Meta-PCA fitted functions in order to calculate disease-specific M-scores for each chaperome functional gene group as described. We then plotted relevant M-scores using polar plots, such that radial axes represent functional processes.

### Physical Protein-Protein Interactome Network Assembly and Curation

Human physical protein - protein interactions (PPIs), hereafter referred to as ‘edges’, were downloaded on 23 Dec 2016 from the BioGRID (Chatr-Aryamontri et al. 2013), IntAct (Kerrien et al. 2012), DIP (Salwinski et al. 2004), and MINT (Licata et al. 2012) databases. In order to obtain a high confidence chaperome physical protein - protein interactome network, we developed a custom Python script to curate raw interactome pairs, or edges, as downloaded from the above databases, considering edges detected by multiple experimental methods as more reliable than those detected by only a single method. Similarly, edges supported by multiple publications are considered at higher confidence than edges supported by only one study. Edges supported by multiple methods and / or multiple studies are collectively referred to as ‘multiple evidence’ (ME), of which those identified by multiple different methodologies represent a subset of highest confidence (MM). The Python script processes the interactome raw data as follows: UniProt IDs are mapped to NCBI Entrez Gene IDs and for each human PPI between any two chaperome members (nodes), interacting partners are mapped to Gene IDs. Only edges annotated with PSI-MI term ‘physical association’ type are considered. Eight different curation levels exist:

|  |  |  |
| --- | --- | --- |
| L1: | *met_single_pub_multiple:* | *1 method (not binary), > 1 PubMed* |
| L2: | *met_single_binary_pub_multiple:* | *1 method (binary), > 1 PubMed* |
| L3: | *met_pub_single_binary:* | *1 method (binary), 1 PubMed* |
| L4: | *met_pub_single:* | *1 method (not binary), 1 PubMed* |
| L5: | *met_pub_multiple:* | *> 1 method (not binary), > 1 PubMed* |
| L6: | *met_multiple:* | *> 1 method (not binary), 1 PubMed* |
| L7: | *binary_met_pub_multiple:* | *> 1 method ($\geq 1$ binary), > 1 PubMed* |
| L8: | *binary_met_multiple:* | *> 1 method ($\geq 1$ binary), 1 PubMed* |

Considering these curation levels, three physical chaperome (CHAP) interactomes of increasing confidence level are obtained (S2 Table):

1. SE-CHAP: 666 unique physical edges between 220 nodes without curation for type or number of evidence (single evidence SE-CHAP)
2. ME-CHAP: 222 unique physical edges between 128 nodes with multiple pieces of evidence detected by ≥ 2 different experimental methods OR reported by ≥ 2 independent studies (PMIDs) (multiple evidence ME-CHAP)
3. MM-CHAP: 132 unique physical edges between 96 nodes detected by ≥ 2 different experimental methods only (multiple method MM-CHAP).

Different PPI source databases may annotate an identical reported PPI to different PSI-MI terms situated at different depth of the same branch within the PSI-MI ontology tree. In these cases, PPIs that are actually only supported by one piece of evidence can unintentionally be mislabelled as multiple evidence PPIs. Our automated quality curation script resolves this problem through up-propagation within the PSI-MI - ontology tree. Assume one PPI is annotated with two different interaction detection methods, A and B, then 1) if PSI-MI ontology tree levels of method A and method B are identical but their PSI-MI terms (IDs) are different, then the methods are considered as different, otherwise A and B are considered the same and the interaction is eliminated from the MM-CHAP interactome, 2) if the level of method A is higher (deeper in the ontology tree) than the level of method B, then the code searches for its parent situated at the same level as method B and compares the parent method ID with B to determine if the methods are identical or different.

### Interactome-Guided 3-D Topographic Maps

We considered 2-dimensional physical interactome information to guide the spatial layout (x-y coordinates) of human chaperome functional ontology families in a 3-dimensional (x-y-z coordinates) topographic representation of chaperome M-score changes between disease and healthy tissue (z coordinate). Physical chaperome protein-protein interactome network data (PPIs) was obtained and curated as described above. We considered a network involving only high quality curated interactions supported by multiple pieces of evidence (ME-CHAP). We used the R package iGraph in order to collapse nodes corresponding to each level 1 functional ontology family into meta-nodes, and edges shared between all members of any two different level 1 functional families into meta-edges, such that meta-node size corresponds to the number of family members and meta-edge thickness represents the number of shared interactions between two families. Meta-node colour is set to reflect gene expression changes of each respective functional family in disease. We then applied a force-directed network graph layout algorithm to the meta-network according to Kamada and Kawai (Kamada and Kawai 1989) and extracted resulting x-y coordinates of each family meta-node in the network. We used Python to draw the meta-network according to the parameters obtained in iGraph to serve as interactome-guided base grid for disease-specific quantitative 3-dimensional topographic network representations. To this end we expanded the 2-dimensional network landscape with Meta-PCA derived chaperome M-score values (z coordinate).

### Proteostasis Profiler (Pro^2^) Web-Tool

We designed a web-based Proteostasis Profiler (Pro^2^) in order to enable visual exploration of the data and results described in this manuscript, obtained through our algorithms and visualisation tools. Pro^2^ is accessible directly at http://www.proteostasys.org or through the resources collection at the Joint Research Center for Computational Biomedicine (JRC-COMBINE) at http://www.combine.rwth-aachen.de/index.php/resources.html. Pro^2^ is implemented using Django (https://www.djangoproject.com/), a web framework written in Python language (https://www.python.org). All the charts in the tool are generated using the plotly platform (https://plot.ly). The Pro^2^ tool itself is hosted on the Heroku platform (https://www.heroku.com) and related code is available at the Github repository at https://github.com/brehmelab/Pro2.

## Acknowledgements

AHE, AS and MB were supported by funding from Bayer AG, and MB is supported by the Joachim Herz Stiftung. The results published here are in part based upon data generated by the TCGA Research Network (http://cancergenome.nih.gov). The authors thank Michael Schubert for providing the processed and voom - transformed TCGA RNA-seq gene expression data and Luz Garcia-Alonso for discussions on protein-protein interaction curation.

### Competing interests

The authors declare no competing financial interests. AAS is a part-time employee of Bayer AG.

### Author contributions

MB conceived, designed and advised overall research. MB and AHE designed the computational analysis strategy. AHE, AS and MB performed computational analyses. AAS and JSR contributed strategic insight. MB wrote the manuscript. All authors reviewed the manuscript.

### Corresponding author

Correspondence to Marc Brehme.

## Supporting information

### Supplementary Figures

**S1 Fig. Correlation of Chaperome GSA and Meta-PCA scores.** Correlation between GSA-scores (Efron and Tibshirani 2007) and Meta-PCA T-statistic values derived using Limma linear modelling on all TCGA cancer groups considered in this study indicates overall correlation (cor = 0.61).

**S2 Fig. Preferential Chaperome Upregulation in Cancers.** Analysis of differential cancer gene expression of chaperome functional subsets, comparing chaperones and co-chaperones as well as ATP-dependent and ATP-independent chaperones (Brehme et al. 2014). See also Fig 3. **A.** Comparing upregulation and downregulation of gene expression of chaperones and co-chaperones using GSA reveals a general upregulation of chaperones and co-chaperones in cancer, with preferential upregulation of chaperones. Colour code indicates chaperone up-regulation of gene expression. Axes represent chaperone downregulation, co-chaperone upregulation and downregulation of gene expression. **B.** Box-and-whisker plots highlight fractions of differentially expressed genes in each chaperome subset for all TCGA cancers assessed, based on A. Differentially expressed genes in each set were obtained by linear modelling (Limma package in R) and considering genes with p value < 0.5 following Benjamini-Hochberg correction. Box boundaries, 25% and 75% quartiles; middle horizontal line, median; whiskers, quartile boundaries for values beyond 1.5 times the interquartile range; small circle, outlier. **C.** Assessing differential expression of ATP-dependent (n = 50) vs. ATP-independent (n = 38) chaperones highlights a preferential upregulation of ATP-dependent chaperones across TCGA cancers **D.** Box plots (drawn as in B.) show fractions of differentially expressed genes in the two sets of ATP-dependent and ATP-independent chaperones for all TCGA cancers assessed, based on C.

**S3 Fig. Compendium of Chaperome Polar Maps.** Chaperome gene expression shifts between healthy and cancer (pp. 1-22) or AD, PD and HD (pp. 23-25) tissue biopsy datasets are quantified by Meta-PCA and visualized in context by plotting resulting M-scores on polar maps as in Fig 6. Blue (healthy) and red (disease) lines represent means across all samples for each disease. Halos represent confidence interval at the 90% quantile range (5% - 95%).

**S4 Fig. Compendium of Chaperome Interactome Meta-Networks.** Chaperome meta-interactome networks for cancers (pp. 1-22) or AD, PD and HD (pp. 23-25) are shown as in Fig 7A. Multiple-evidence chaperome (ME-CHAP) edges and nodes are collapsed onto meta-edges and meta-nodes. Node size and edge thickness correspond to the number of functional family member nodes and the sum of the number of inter-family edges, respectively. Node colour indicates combined disease gene expression changes quantified via Meta-PCA.

**S5 Fig. Compendium of Chaperome 3-D Topographic Maps.** 3D topographic maps of cancer (pp. 1-22) or AD, PD and HD (pp. 23-25) chaperome alterations are obtained by projecting gene expression changes between disease and healthy counterpart biopsy gene expression (z dimension) onto the high-confidence chaperome meta-interactome (ME-CHAP) graph layout (x-y dimensions).

### Supplementary Tables

**S1 Table. Human Chaperome and Proteasome Functional Gene Ontology. Tab S1A.** List of 332 human chaperome genes, expert curated by functional ontology groups at six levels of increasing detail (Level 1 = broad >> level 6 = detailed). For each entry, HGNC gene symbol, EntrezID, and functional ontology annotation levels are indicated. Human chaperome as in (Brehme et al. 2014), **Tab S1B.** List of 43 proteasome genes. For each entry, HGNC gene symbol, EntrezID, and functional ontology annotation levels are indicated. Proteasome complex annotation according to HGNC (Gray et al. 2015).

**S2 Table. Human Chaperome Physical Protein-Protein Interactions.** Human physical protein-protein interactions (PPIs) were obtained for 332 human chaperome members from curated public databases reporting human PPIs with PSI-MI method annotations (IntAct, BioGrid, MINT, DIP). For each node, Entrez Gene ID and corresponding HGNC symbol, chaperome functional ontology annotation levels 1 and 2, interaction detection method MI code as annotated in the respective source database, and unique methods per PPI following curation, curation evidence level, PMIDs with corresponding literature evidence for each interaction, source databases, and respective MI method IDs are indicated. **Tab S2A**. 666 curated physical human protein - protein interactions between 220 chaperome nodes supported by any number of evidence, including interactions reported only by one single study (PMID). **Tab S2B**. 222 curated high-confidence physical human protein - protein interactions between 128 chaperome nodes supported by multiple pieces of evidence, including interactions reported by multiple methods or studies (PMIDs). **Tab S2C**. 132 curated high-confidence physical human protein - protein interactions between 96 chaperome nodes supported by multiple pieces of evidence, including only interactions reported by multiple methods.

### Other Supplementary Resources

All R and Python scripts and code related to this manuscript are accessible through the proteostasis profiler (Pro^2^) Github repository at https://github.com/brehmelab/Pro2.

